# A Highly Conserved ABC Transporter Mediates Cello-Oligosaccharide Uptake in the Extremely Thermophilic, Lignocellulolytic Bacterium *Anaerocellum (f. Caldicellulosiruptor) bescii*

**DOI:** 10.1101/2025.02.26.640372

**Authors:** Hansen Tjo, Virginia Jiang, Anherutowa Calvo, Jerelle A. Joseph, Jonathan M. Conway

## Abstract

Cellulose deconstruction and utilization are key to unlocking renewable fuel and chemical production. *Anaerocellum bescii* (formerly *Caldicellulosiruptor bescii*) is an extremely thermophilic cellulolytic bacterium, among the most effective at degrading lignocellulosic biomass due to its arsenal of multi-domain cellulases and hemicellulases. However, little is known about how it transports the assorted sugars released from lignocellulose degradation into the cell for catabolism. Among its twenty-three ATP-Binding Cassette (ABC) sugar transporters, the mechanism for uptake of cello-oligosaccharides released from cellulose degradation remains unclear. Here, we identify an ABC transporter locus (*Athe_0595 — 0598*), highly conserved in the genus with two extracellular binding proteins, Athe_0597 and Athe_0598. Biophysical analyses, including Differential Scanning Calorimetry (DSC) and Isothermal Titration Calorimetry (ITC), reveal that Athe_0597, binds cello-oligosaccharides of varying lengths (G2-5), while Athe_0598 is specific to cellobiose (G2). Computational modeling of ligand docking supports these findings and sheds light on the subsite configuration of the substrate binding proteins. To assess its physiological importance, we genetically deleted this transporter locus in *A. bescii* strain HTAB187, which does not grow on cellulose and grows poorly on cellobiose. Comparison of growth with a *msmK* deletion strain that cannot consume oligosaccharides shows that HTAB187 can grow on non-cello-oligosaccharides (e.g. maltose) or monosaccharides. Taken together, this study integrates biophysical characterization, structural modeling, and genetic perturbation to elucidate how *A. bescii* transports cello-oligosaccharides release from cellulose, opening doors for its future use in applied bioprocessing contexts.

**Importance:** *Anaerocellum bescii* is the most thermophilic cellulolytic bacterium known, and holds potential for bioprocessing lignocellulosic biomass into renewable fuels. Its diverse ATP-Binding Cassette (ABC) sugar transporters make it a valuable model for studying thermophilic sugar uptake. Here, we identify a single ABC transporter with two substrate binding proteins (Athe_0597 and Athe_0598) responsible for cello-oligosaccharide uptake. Genetic deletion of this transporter impaired growth on cellobiose and eliminated growth on cellulose. This is the first genetic manipulation in *A. bescii* to modulate transport of a specific sugar. We also characterize the substrate specificity of the substrate binding proteins associated with the locus; one binds various cellodextrins (G2-5), while the other specifically binds cellobiose (G2). Computational modeling reveals how each sugar docks within the binding pocket of these proteins. Understanding the mechanism of cello-oligosaccharide uptake by *A. bescii* expands opportunities for its metabolic engineering and furthers our understanding of thermophilic sugar transport.

## Introduction and Background

Confronted by an increasingly warming climate, society must replace petroleum-based supply chains with renewable bio-based fuels and chemicals derived from lignocellulosic feedstocks (1,2). Yet for decades scientific progress has been stymied by the physical and chemical recalcitrance of lignocellulose, impeding deconstruction and conversion at the large scales needed to compete with cheap petroleum (3,4). Recent genetic engineering efforts make lignocellulolytic thermophiles of interest for widespread use in lignocellulose bioprocessing (5–7). Realizing this prospect demands a more comprehensive understanding of thermophilic physiology, particularly concerning sugar uptake (8,9).

The extremely thermophilic, lignocellulose-degrading bacterium *Anaerocellum bescii* is the most thermophilic cellulolytic bacterium known with an optimal growth temperature of 75°C (10–12). It possesses vital attributes for consolidated bioprocessing of lignocellulosic feedstocks: efficiently degrading cellulose and hemicellulose into oligosaccharides, while simultaneously metabolizing pentose and hexose sugars (13). Its unique Glucan-Degradation-Locus (GDL) expresses high levels of multi-domain glycoside hydrolases with catalytic activity on cellulose, xylans, and mannans (14,15). *A. bescii* is also armed with a rich secretome with many additional catalytic and non-catalytic proteins, e.g., glycoside hydrolases, substrate-binding proteins, acting in concert to maximize the degradation and cellular uptake of the diverse polysaccharides that make up lignocellulose, e.g., cellulose, xylans, arabinoxylans, pectin. Its secretome composition is also known to adapt to external substrates, underscoring the complexity of its sugar degradation and transport mechanisms that also evidences its efficient adaptation to carbohydrate-limiting environments (15–17). And, multiple genetic tools including methods for chromosomal modification, an antibiotic-based selection marker, as well as xylose-inducible promoter, have been developed over the past decade (18,19). These tools have enabled metabolic engineering in *A. bescii* to produce a wide range of valuable products including ethanol, acetone, and 2,3-butanediol (2,3-BDO) at elevated yields (6,20,21). Despite these advances, however, relatively little is understood about how *A. bescii* imports the diverse range of sugars released from lignocellulose into the cell for metabolism (22).

*A. bescii* primarily utilizes ATP-Binding Cassette (ABC) sugar transporters that hydrolyze two molecules of ATP for the translocation of one sugar molecule into the cytoplasm (23,24). Twenty-three such ABC transporters have been annotated in the bacterium, with the twenty-fourth sugar transporter being a Phosphotransferase System (PTS) transporter for putative fructose transport (13,25). ATP-Binding Cassette transporters comprise a hydrophobic trans-membrane protein domain, a cytoplasmic ATPase domain, and an extracellular substrate-binding domain (24). This substrate-binding domain, otherwise referred to as substrate-binding protein (SBP), helps indicate the sugar specificity of its associated transporter as it is responsible for binding extracellular sugars and bringing them to the trans-membrane domain (22,26,27). In some cases, however, the SBP can bind to substrates that are too large in size for the transmembrane domain to move across the membrane (28,29). There is also genetic evidence to suggest that MsmK, a promiscuous ATPase encoded by gene locus Athe_1803, is responsible for powering all oligosaccharide ABC sugar transporters in *A. bescii* (13). Though a number of transcriptomic studies have yielded predictions of the specificity of the ABC transporters in *A. bescii,* only the two transporters for the maltodextrin system have been experimentally validated thus far (13,25). There have not been studies that validate how cellulose-derived oligosaccharides, on which *A. bescii* grows to its highest cell densities, are transported into the cell (16).

In this study, we report the first ever molecular investigation of cello-oligosaccharide transport in *A. bescii.* We show that two substrate-binding proteins, Athe_0597 and Athe_0598, are responsible for utilization of cello-oligosaccharides in *A. bescii*. We demonstrate that these proteins bind cello-oligosaccharides at characteristically high binding affinities, with *in silico* models that structurally elucidate how these sugars are bound. By genetically deleting the cello-oligosaccharide ABC transporter locus in *A. bescii,* we establish its role in cello-oligosaccharide transport *in vivo*. This deletion strain exhibited reduced growth on the disaccharide cellobiose and was entirely unable to grow on cellulose, demonstrating the critical role of these transporters in cellulose utilization. These insights pave the way for engineering *A. bescii* strains with novel, customized carbohydrate utilization pathways.

## Results

### Identification of the cello-oligosaccharide ABC Transport System in *A. bescii*

*A. bescii* contains one loci encoding an ABC transporter (Athe_0595–0598) predicted to mediate cello-oligosaccharide transport (Figure 1A) based on transcriptomic work by VanFossen et al. (2009) in *Caldicellulosiruptor saccharolyticus* and Rodionov et al. (2021) in *A. bescii* (18,19). Rodionov et al. (2021) predicted that Athe_0597 and Athe_0598 encode for ABC substrate-binding proteins (13). Notably, their expression was up-regulated when *A. bescii* was grown on cellulose (13). Yet both transporter substrate predictions from Vanfossen et al. (2009) and Rodionov et al. (2021) to this date have not been experimentally validated.

**Figure 1:**
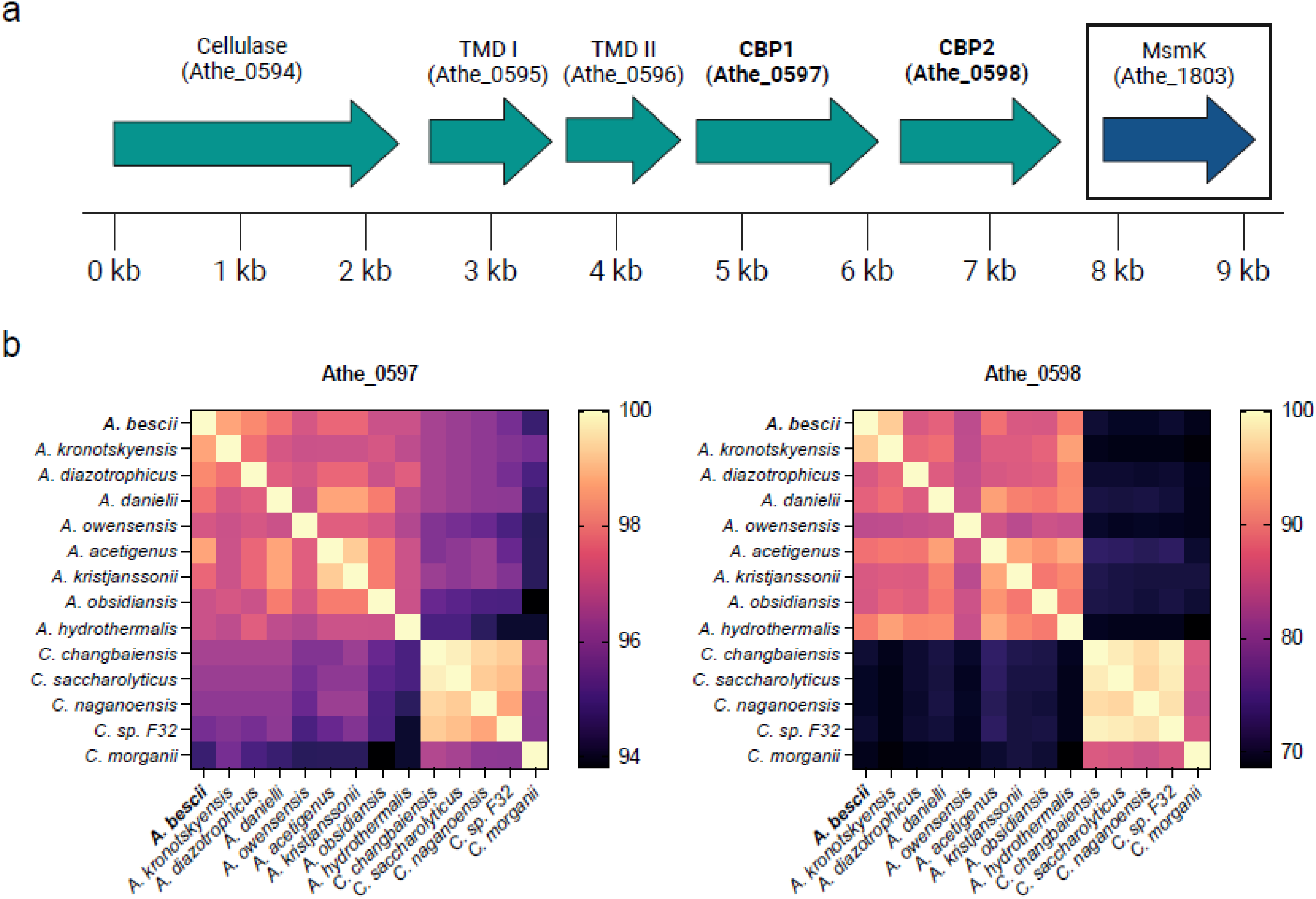
A) Genomic organization of cello-oligosaccharide ATP-binding cassette (ABC) transporters and neighboring genes associated with cellulose utilization in *Anaerocellum bescii.* B) BLAST analysis of protein sequence conservation of cello-oligosaccharide-binding proteins Athe_0597 and Athe_0598 across the *Anaerocellum* and *Caldicellulosiruptor* genus.

The genomic context of Athe_0597 and Athe_0598 suggests that they are involved in cello-oligosaccharide uptake. Both genes are co-localized with Athe_0594, which encodes an extracellular GH5-CBM28 endoglucanase that degrades cellulose into glucose and cello-oligosaccharide products (14,15,30). This operon was found to be up-regulated over 10-fold in *A. bescii* strains grown on cellulose and cellobiose growth substrates (13). Interestingly, both cello-oligosaccharide-binding proteins (CBPs) Athe_0597 and Athe_0598 may share the same transmembrane domains Athe_0595– 0596 as no other transmembrane domains are proximal. No ATPase gene is also present; deletion of the promiscuous *msmK* ATPase (Athe_1803) was found to deactivate all oligosaccharide transporters and consequently inhibit growth on cello-oligosaccharides (13). This suggests that the cello-oligosaccharide ABC transporter as encoded by the Athe_0595–0598 locus is powered by the promiscuous *msmK* ATPase (Athe_1803) (13). The putative ABC cello-oligosaccharide transporter locus appears widely conserved across the *Anaerocellum* and *Caldicellulosiruptor* genera, though to a lesser extent in the latter (Figure 1B) (the *Anaerocellum* genus was recently split from the *Caldicellulosiruptor* genus) (12). All *Anaerocellum* and *Caldicellulosiruptor* species possess genes encoding the heterodimeric trans-membrane domain, as well as genes encoding for two separate cello-oligosaccharide-binding proteins. Athe_0597 appears much more highly conserved across both genera; the *C. morganii* homolog, possessing the highest sequence deviation from the *A. bescii* version, still retains a relatively high amino acid sequence identity of 94.8%. Conversely, Athe_0598 homologs exhibit higher sequence variability, particularly amongst *Caldicellulosiruptor* species as in the cases of *C. saccharolyticus, C. changbaiensis, C. naganoensis, C. sp. F32,* and *C. morganii.* Yet even within the *Anaerocellum* genus, *A. owensensis* has the lowest amino acid sequence identity at 83.6%.

### Biophysical Determination of cello-oligosaccharide Specificity

Differential scanning calorimetry (DSC) is useful for screening the carbohydrate specificity of extremely thermophilic substrate-binding proteins (22,31). As it can reach temperatures as high as 130°C, it can denature *C. bescii* substrate-binding proteins that typically possess native melting temperatures above 92°C, the limit of traditional circular dichroism (CD) methods (22). In DSC, a constant rate of heating raises the heat capacity of the protein until it fully denatures. The temperature corresponding to the maximum heat capacity of the protein, implying complete denaturation, is the native melting temperature *T*_*m*_ of the substrate-binding protein in its *apo* state. When bound to its cognate substrates, however, these substrate-binding proteins acquire a closed conformation with higher thermal stability – a process referred to as the “Venus-flytrap” mechanism (22,26). The difference in melting temperatures across the *apo* and *holo* states, defined as |𝛥*T*_*m*_| = *T*_*m*,*holo*_ − *T*_*m,apo*_, denotes the sugar specificity of a given substrate-binding protein; sugars with higher affinity will yield greater 𝛥*T*_*m*_ values (22).

DSC melting curves show that Athe_0597 possesses a strong affinity for cello-oligosaccharide substrates (G2–G5). In its *apo* state, Athe_0597 exhibits a melting curve with a lower peak at *T*_*m*,1_ = 77.6 °C and a higher peak at *T*_*m*,2_ = 91.5 °C (Figure 2A). The higher peak at *T*_*m*,2_ = 91.5 °C is the presumptive melting temperature, but it is possible that in the absence of a bound ligand a subunit begins denaturing at *T*_*m*,1_ = 77.6 °C. The melting curve of Athe_0597 collapses into a single peak in the presence of cognate sugars cellobiose (G2), cellotriose (G3), cellotetraose (G4), and cellopentaose (G5) with increased melting temperatures of 98.4°C to 100.8°C. Curiously, melting temperature shifts are nearly identical for all tested cello-oligosaccharides (𝛥*T*_*m*_ > 11°C). The rank ordering of melting temperature increases also do not neatly correlate with oligosaccharide size. This contrasts the larger differential 𝛥*T*_*m*_ values observed with the binding of *A. bescii* maltodextrin-binding protein Athe_2574 across maltodextrin ligands such as maltose (𝛥*T*_*m*_ ∼ 5.03°C) to maltoheptaose (𝛥*T*_*m*_ > 12°C) (22). Athe_0597 also shows no changes in its melting profile relative to its *apo* state in the presence of glucose, demonstrating its lack of affinity for the monosaccharide.

**Figure 2:**
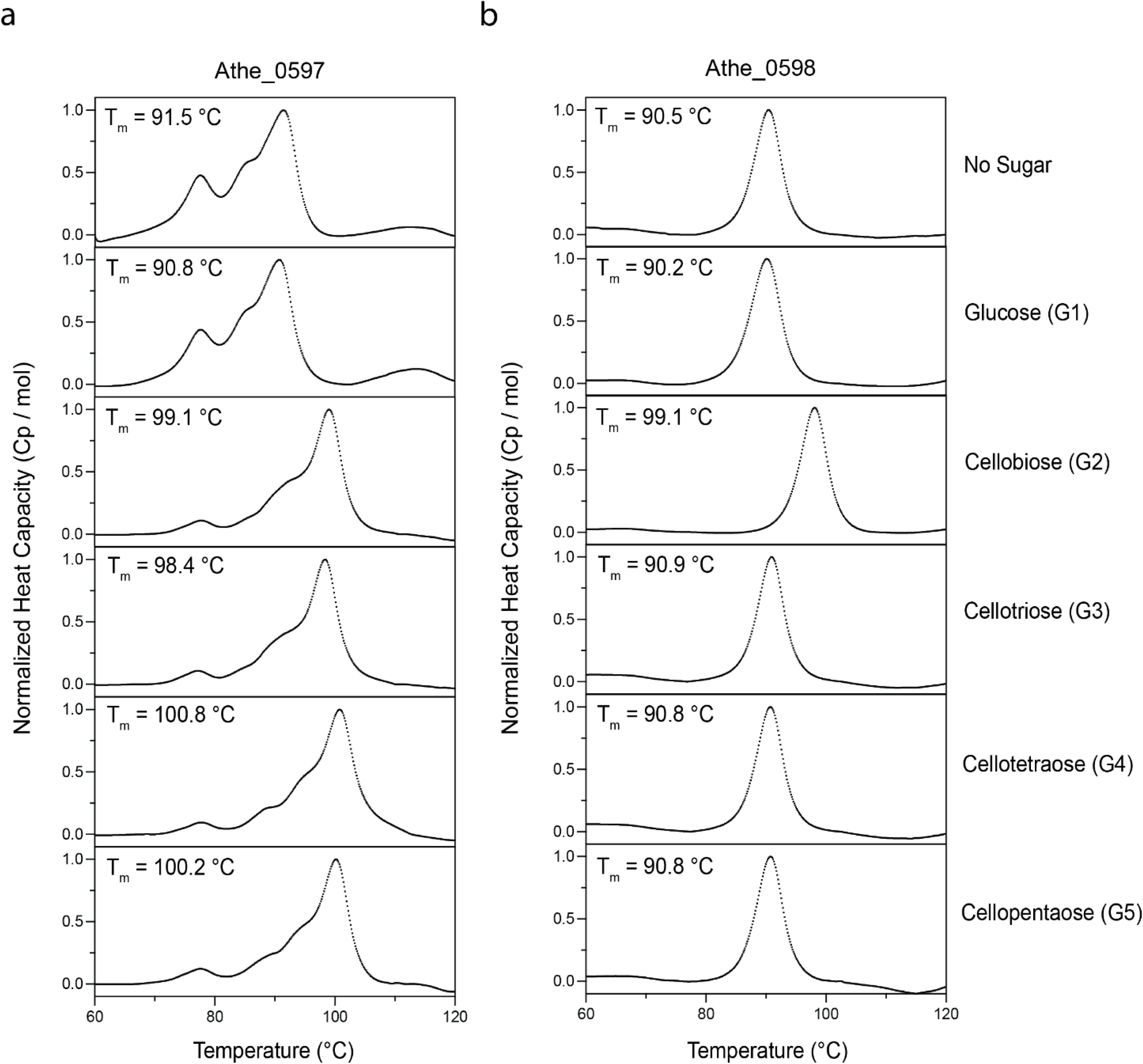
Normalized differential scanning calorimetry (DSC) screens of A) Athe_0597 and B) Athe_0598 mixed with glucose as well as 𝛽-linked cello-oligosaccharides of different lengths: cellobiose (G2), cellotriose (G3), cellotetraose (G4), cellopentaose (G5). Athe_0597 is shown to bind all tested cello-oligosaccharides from cellobiose to cellopentaose, whereas Athe_0598 only binds cellobiose. All DSC screens were performed at a temperature range of 60 - 130°C, in 50 mM HEPES and 300 mM NaCl pH 7.0 buffer.

Athe_0598 shows uniform melting even in its unliganded *apo* state (Figure 2B). The melting curve profile in its unliganded *apo* state is similar to when it is bound to its cognate substrate cellobiose (G2). Indeed, Athe_0598 only exhibits affinity towards cellobiose as a substrate. Larger cello-oligosaccharide substrates from cellotriose to cellopentaose, as well as the monomer unit glucose, did not result in any *T*_*m*_ shifts, suggesting none of these oligosaccharides are able to bind Athe_0598.

Next, ITC experiments were performed to probe the binding interactions between the CBPs and their cognate cello-oligosaccharide ligands. Figure 3 illustrates the binding isotherms and isothermal titration curves for all sugar–protein pairs that were found to bind according to the DSC, with key measurements including dissociation constants 𝐾_𝑑_ and stoichiometry (𝑛) summarized in Table 1. ITC data reinforced results from the DSC, showing that Athe_0597 indeed binds cellobiose, cellotriose, cellotetraose, and cellopentaose with micromolar dissociation constants typical of high affinity substrates in ABC sugar transporter systems (32,33). While 𝐾_𝑑_ values at first glance appear relatively similar for all cello-oligosaccharide substrates, upon closer inspection of the association constant 𝐾_𝑎_ values, it becomes clear that Athe_0597 has the highest affinity for cellobiose (Table 1). The 𝐾_𝑎_ for cellobiose is on order-of-magnitude higher than for all other cello-oligosaccharides, including cellotriose when taking into account the observed error (Table 1). It is worth noting that the enthalpies of binding for Athe_0597 with all cello-oligosaccharides are endothermic, which while not unique, is less commonly observed for ABC sugar-binding proteins (32,33). The binding of both Athe_0597 and Athe_0598 to cello-oligosaccharide substrates also appear to be consistently entropically driven, with *T*𝛥*S* values significantly exceeding 𝛥𝐻 values in all measured CBP-cello-oligosaccharide combinations (Table 1).

**Figure 3:**
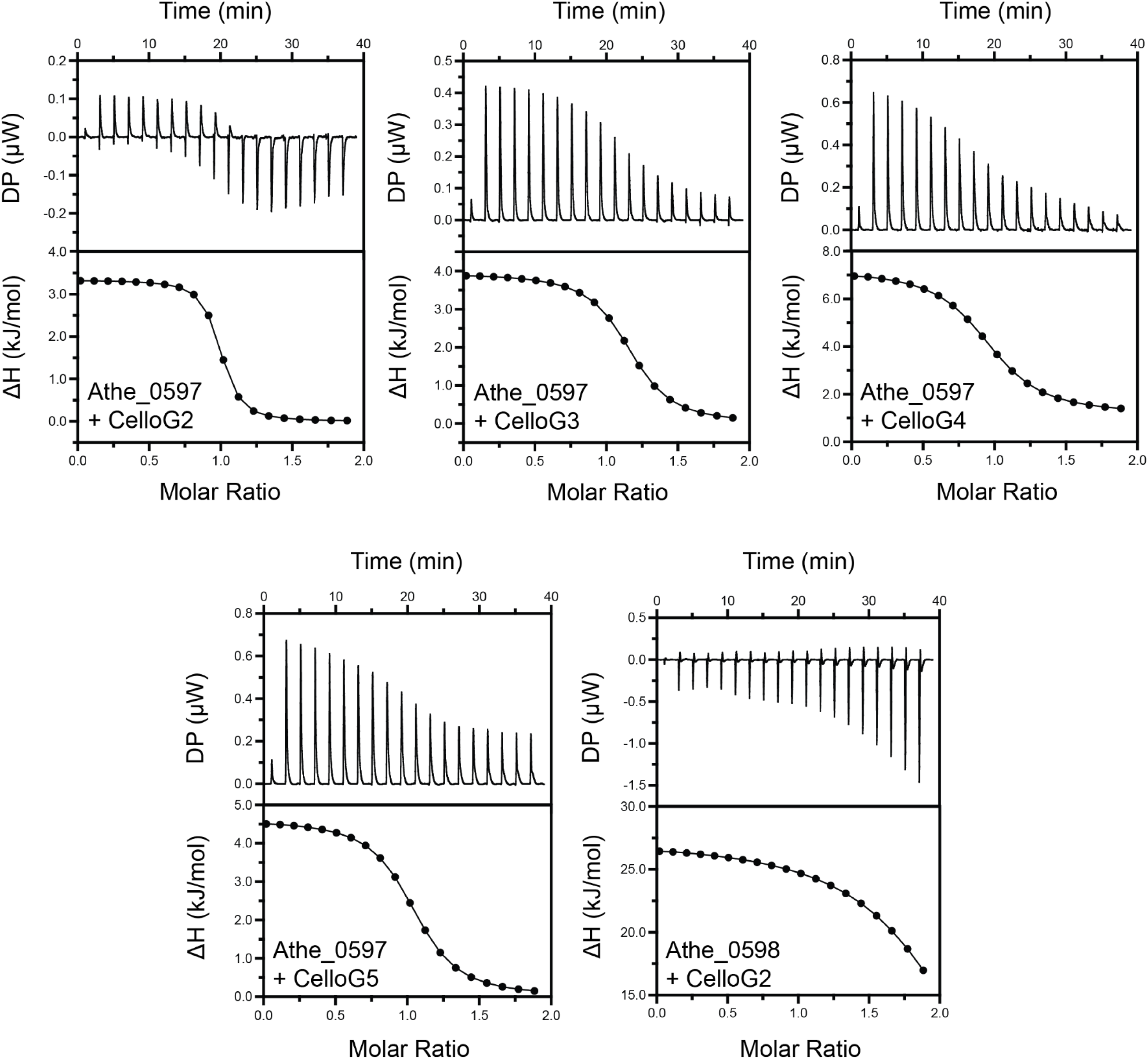
Representative isothermal titration calorimetry (ITC) screens of all CBP and cello-oligosaccharide pairings that were successful from DSC in Figure 2. These combinations include Athe_0597 with cellobiose (G2), cellotriose (G3), cellotetraose (G4), and cellopentaose (G5), as well as Athe_0598 with cellobiose (G2). Both raw isothermal titration curves as well as integrated binding isotherms are shown for each CBP and cello-oligosaccharide pairing.

**Table 1:** Biophysical and thermodynamic parameters of binding between CBPs and - linked cellodextrins as determined by DSC and ITC. CBP-sugar combinations denoted with a “-” were not tested via the ITC as DSC measurements indicated either weak or absent binding.

| Protein | Sugar | $T_m$ (°C) | $\Delta T_m$ (°C) | $n$ | $K_d$ (μM) | $K_a$ ($\times 10^5 \text{ M}^{-1}$) | $T$ (°C) | $\Delta H$ ($\frac{\text{kcal}}{\text{mol}}$) | $T \times \Delta S$ ($\frac{\text{kcal}}{\text{mol}}$) | $\Delta G$ ($\frac{\text{kcal}}{\text{mol}}$) |
| --- | --- | --- | --- | --- | --- | --- | --- | --- | --- | --- |
| Athe_0597 | Glucose | 90.79 | -0.7 | - | - | - | - | - | - | - |
| | CelloG2 | 99.06 | 7.57 | 0.95<br>$\pm 0.02$ | 0.228<br>$\pm 0.10$ | 43.86 | 25 | 3.33<br>$\pm 0.13$ | -12.4 | -9.07 |
| | CelloG3 | 98.38 | 6.89 | 1.13<br>$\pm 0.01$ | 0.942<br>$\pm 0.12$ | 10.62 | 25 | 3.94<br>$\pm 0.08$ | -12.2 | -8.22 |
| | CelloG4 | 100.79 | 9.30 | 0.95<br>$\pm 0.03$ | 1.97<br>$\pm 0.52$ | 5.08 | 25 | 6.04<br>$\pm 0.32$ | -13.8 | -7.79 |
| | CelloG5 | 100.21 | 8.72 | 1.02<br>$\pm 0.08$ | 1.18<br>$\pm 0.12$ | 8.47 | 25 | 4.61<br>$\pm 0.08$ | -12.7 | -8.09 |
| Athe_0598 | Glucose | 90.17 | -0.36 | - | - | - | - | - | - | - |
| | CelloG2 | 99.05 | 8.53 | 2.13<br>$\pm 0.24$ | 2.84<br>$\pm 0.36$ | 3.52 | 25 | 27.2<br>$\pm 10.9$ | -34.7 | -7.57 |
|  | CelloG3 | 90.91 | 0.39 | - | - | - | - | - | - | - |
|  | CelloG4 | 90.81 | 0.281 | - | - | - | - | - | - | - |
|  | CelloG5 | 90.79 | 0.26 | - | - | - | - | - | - | - |

### Docking Simulations Elucidate Structural Context to Cello-oligosaccharide Binding

Computational models of binding affinity show that Athe_0597 is able to bind cellobiose, cellotriose, cellotetraose, and cellopentaose while Athe_0598 primarily favors binding to cellobiose (Figure 4C) (Table S1). These conclusions largely match results obtained through experiments. The modeled Athe_0598 likely retains some affinity for cellotriose as the closest-match homologs with crystal structures that were used as templates for the model have affinity for larger sugars as well as cellobiose (34). Simulations reveal that each CBP possesses a ligand binding pocket composed of subsites where glucosyl monomers can be docked (Figure 4). Favorable binding free energies are driven by entropic contributions such as the release of water molecules from the protein binding site and the formation of specific enthalpic interactions between each glucosyl monomer and the protein at each subsite.

**Figure 4.**
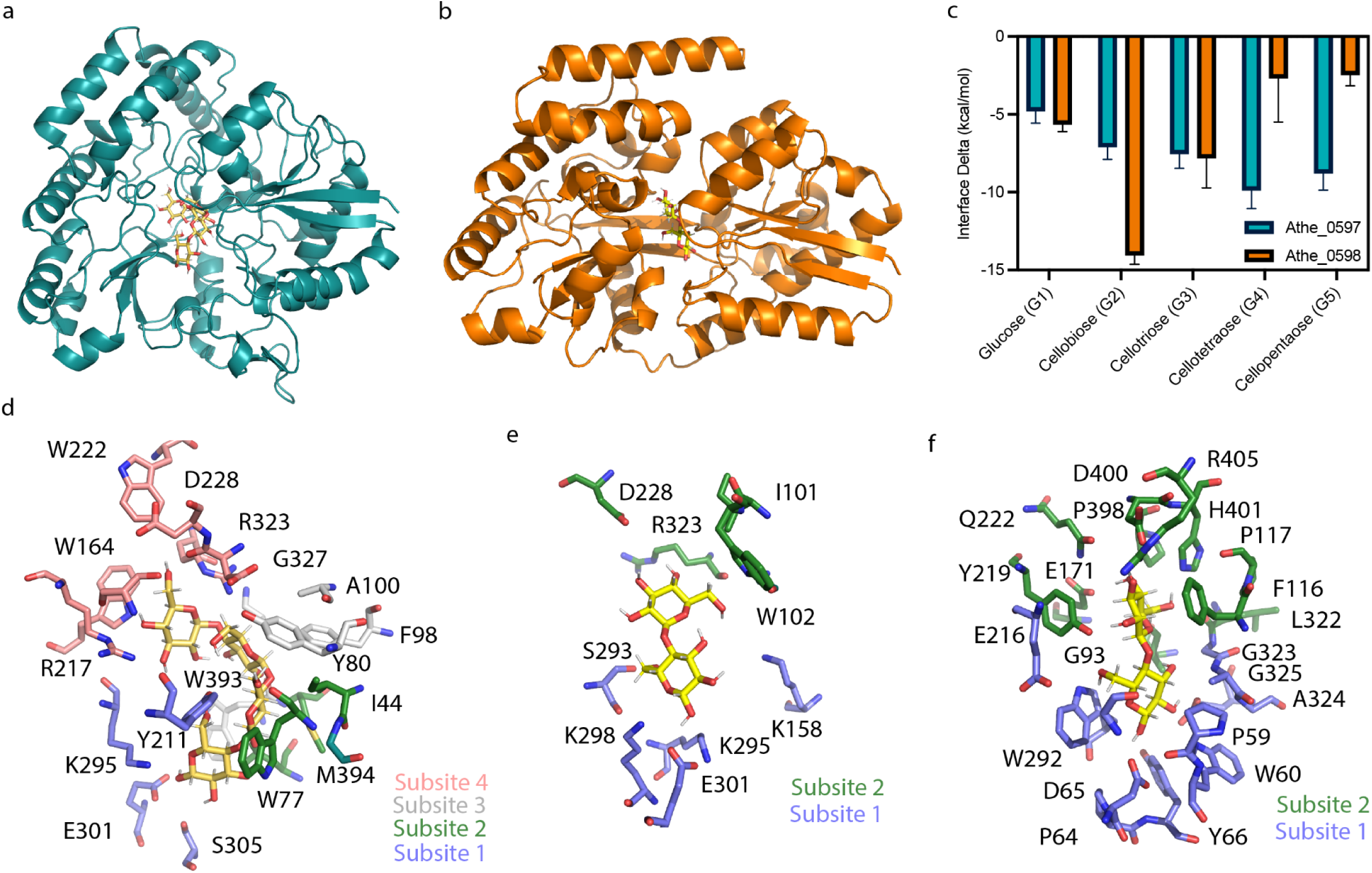
A) Computational model of Athe_0597 as visualized on PyMol. B) Computational model of Athe_0598 as visualized on PyMol. C) Interface energy deltas, as a proxy for free energies of binding, are shown for each simulated combination of cello-oligosaccharide and CBP. The rank-order affinity of Athe_0597 and Athe_0598 with their respective cello-oligosaccharide substrates are aligned with experimental results D) Binding of the highest affinity ligand to Athe_0597, cellotetraose (G4), is mediated by multiple residues across four subsites in its binding pocket. E) Only subsites 1 and 2 are used to coordinate binding from Athe_0597 to its cognate disaccharide ligand cellobiose (G2). F) Multiple residues in the binding pocket of Athe_0598 mediate the two subsites that coordinate its primary cognate ligand, cellobiose (G2).

The rank-ordering for substrate binding affinity as measured by ligand docking for Athe_0597 aligns with the rank-ordering by DSC, where cellotetraose is the most favored (Figure 4C). Although ITC and DSC results show a different rank-ordering of binding affinity, the larger errors associated with 𝐾_𝑑_ values from the ITC, and the fact that the DSC is a direct measure of added thermal stability from ligand binding, impels us to place more weight on the DSC’s rank-ordering of substrate affinity. The reducing end of the cellotetraose is bound by subsite 1 with hydrogen bonds between K296, E301, and S305. An internal glucosyl monomer 2 would occupy subsite 2, while a third monomer occupies subsite 3, forming a hydrogen bond with Y80. The nonreducing glucosyl cap occupies subsite 4, with hydrogen bonds between R217 and R323. As shown through ITC, entropic contributions primarily drive substrate affinity (Table 1). Each subsite contains hydrophobic amino acids: Y211 in subsite 1, I44, M394, W77 in subsite 2, A100, F98, W393 in subsite 3, W222 and G327 in subsite 4 (Figure 4D). In the apo case, these residues would remain solvent-accessible. Upon ligand binding, the cello-oligosaccharide displaces these bound water molecules and the release of water to the bulk solvent leads to a net increase in solvent translational entropy.

In the Athe_0598 case, there are fewer available subsites for forming enthalpic interactions and fewer hydrophobic residues in the binding pocket that contribute to desolvation entropy. Athe_0598 had the most favorable binding interface energy with the disaccharide cellobiose, as only two subsites are available (Figure 4F). The reducing end occupies subsite 1, where E167, Y219, and D64 form hydrogen bonds and W60, Y66, and W292 form hydrophobic contacts. The nonreducing end is bound by subsite 2, where D400, H401, and E171 form hydrogen bonds and I169, L322, P117, P398 and F116 form hydrophobic contacts (Figure 4F). These computational results suggest a possible mechanism for the entropically driven binding of various cello-oligosaccharides to Athe_0597 and Athe_0598. Additionally, the selective binding of cellobiose to Athe_0598 may be due to the availability of subsites that could form possible enthalpic interactions, or lack of available hydrophobic patches for entropic desolvation.

### Genetic Deletion of the cello-oligosaccharide ABC Transporter (Athe_0595**–** Athe_0598) Disrupts Growth on Cellobiose and Cellulose

To probe its function in carbohydrate assimilation, the ABC cello-oligosaccharide transporter locus (Athe_0595–0598) was deleted *in vivo* via homologous recombination (19) from the uracil biosynthesis-deficient parent strain MACB1018 (Δ*pyrE*). The cello-oligosaccharide transporter deletion strain HTAB187 (Δ*pyrE* Δ*athe_0595-0598)* was generated using maltose as the carbon source following transformations.

We conducted growth curves using strain HTAB187 on individual carbon sources including glucose, maltose, cellobiose, and microcrystalline cellulose (Avicel) (Figure 5). We also monitored growth curves for the parent strain MACB1018 (Δ*pyrE*) and the MsmK deletion strain MACB1080 (Δ*pyrE* Δ*msmK*). For the latter, we sought to observe whether a single cello-oligosaccharide transporter deletion results in similar growth disruption compared to inactivation of all oligosaccharide transporters in *A. bescii* (13). Unsurprisingly, all three strains demonstrated similar growth behavior on glucose: HTAB187 behaves similarly to MACB1018, while MACB1080 exhibits a slightly lower growth rate compared to MACB1018 during the exponential phase that is consistent with previous results (13). HTAB187 also exhibited no deviations in growth behavior compared to MACB1018 on maltose, while growth on the disaccharide is impaired for MACB1080 due to inactivation of its maltodextrin transporters through loss of ATPase MsmK. On cellobiose, all three strains displayed distinct growth behaviors. Growth on cellobiose was clearly impaired for HTAB187 compared to MACB1018, though the defect was not significant enough to prevent growth entirely. Surprisingly, MACB1080 showed no growth whatsoever on cellobiose in contrast with literature results that suggest minimal growth on the disaccharide (13). Despite some growth on cellobiose, HTAB187 appears to be entirely incapable of growth on cellulose; MACB1080 also showed no growth on cellulose, consistent with prior reports (13). Visual inspection of the turbidity of cultures from all three strains on cellulose suggested that only MACB1018 was thriving (Figure S3). Broadly, deletion of the Athe_0595–0598 cello-oligosaccharide transporter locus results in 1) significant, though not complete, impairment of growth on cellobiose, and 2) elimination of growth on cellulose, without apparent defects in the uptake of other glucose-based substrates such as glucose or maltose. Although it is possible that cellobiose can be extracellularly hydrolyzed into glucose, which HTAB187 can consume, it is more likely that low affinity transport through another ABC transporter explains the limited growth on cellobiose. If the former was the case, we would have expected partial MACB1080 growth on cellobiose given that it can still grow on glucose. Overall, these results indicate that Athe_0598–0598 is the major cello-oligosaccharide transporter in *A. bescii*.

**Figure 5.**
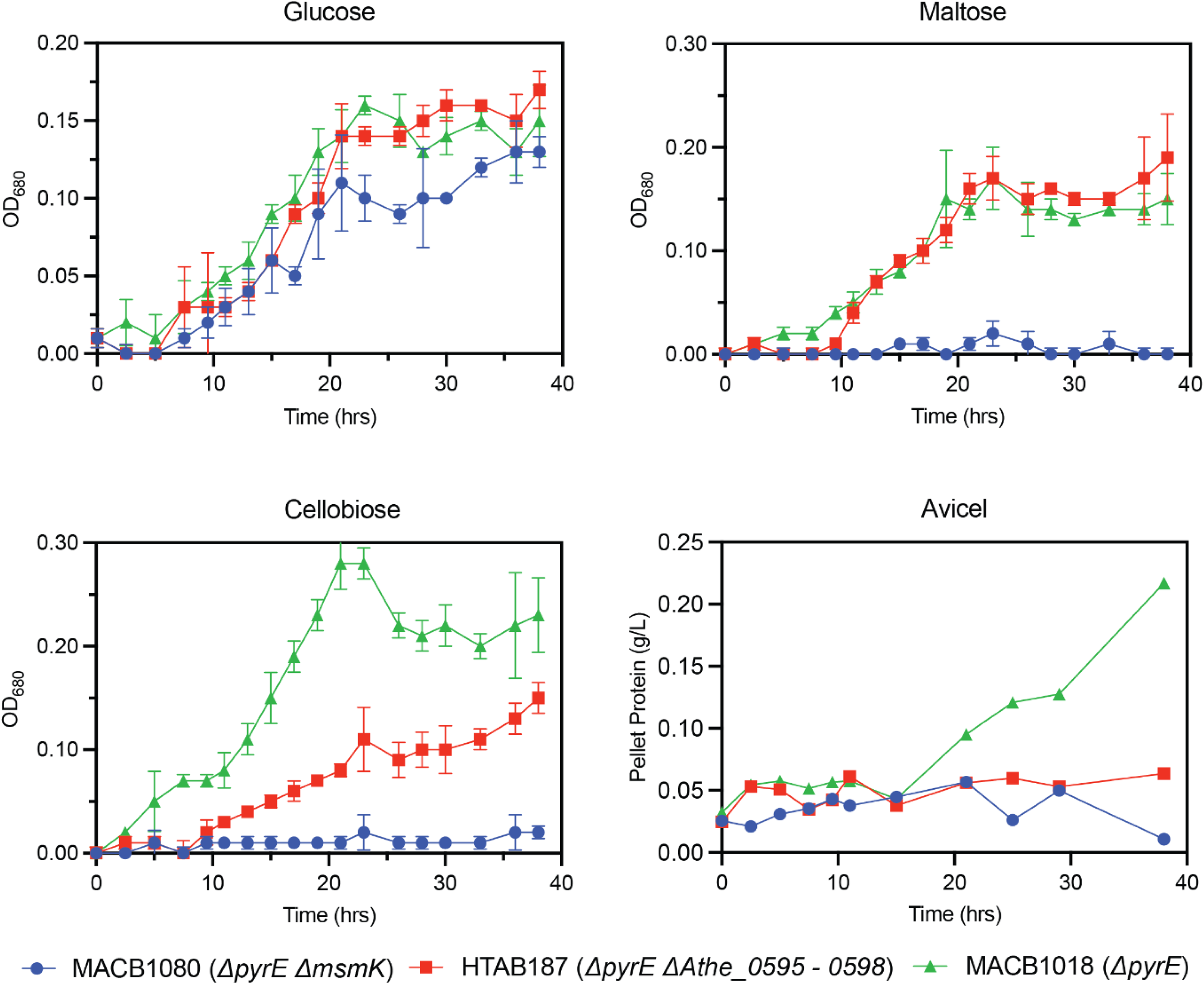
Growth curves of axenic, engineered *A. bescii* strains MACB1018 (green triangles), MACB1080 (blue circles), and HTAB187 (red squares) on carbohydrate substrates glucose, maltose, cellobiose, and microcrystalline cellulose (Avicel). All growth curves were performed in triplicate. For growth curves on glucose, maltose, and cellobiose, each data point represents the mean Optical Density at 680 *nm* (*OD*_680_) value. Error bars denote the standard deviation. Growth curves on Avicel were performed in triplicate, but time point samples are mixed prior to pellet protein quantitation as done previously (44).

## Discussion

Here using biophysics, structural analysis, and genetic knockouts we show that Athe_0595–Athe_0598 is the primary ABC transporter that enables *A. bescii* to utilize cello-oligosaccharides. Through differential scanning calorimetry, we screened the substrate specificity of putative cello-oligosaccharide-binding proteins Athe_0597 and Athe_0598, leveraging the fact that substrate-binding proteins adopt a more thermostable conformation when bound to their cognate substrates. We further characterized and quantified the thermodynamics of cello-oligosaccharide binding using isothermal titration calorimetry, showing that both CBPs bind to β-glucan substrates with dissociation constants in the μM range characteristic of high affinity ABC sugar transporters. Through sequence and structural alignment to known thermophilic orthologs, we further elucidated how such cello-oligosaccharide substrates are docked within their respective binding pockets. Finally, we complemented *in vitro* and *in silico* characterization of cello-oligosaccharide transport with knockout of the Athe_0595–0598 locus *in vivo,* and subsequently study of changes to growth behavior on multiple carbohydrate substrates. We show that deletion of this transporter is necessary and sufficient for eliminating growth on cellulose, and greatly impairs growth on cellobiose, highlighting the principal role of this locus in cello-oligosaccharide uptake.

Broadly, both ITC and DSC corroborated results from one another in showing that Athe_0597 and Athe_0598 both bind to cello-oligosaccharide substrates, and that bound substrates correspond to 𝐾_𝑑_ constants in the uM range as well as positive *T*_*m*_ shifts, relative to the absence of ligand, consistent with experimental study of other thermophilic substrate-binding proteins (Table 1) (22,32,33). Although Athe_0598 binds to cello-oligosaccharides of various length from cellobiose to cellopentaose with similar *T*_*m*_ shifts, Athe_0597 only binds to cellobiose. While ITC and DSC largely agree on the cello-oligosaccharides that induce high affinity binding with Athe_0597, differences in thermal stabilization and micromolar range are not as strictly correlated. We found a decreasing rank order between *T*_*m*_ shift and oligosaccharide length with Athe_0597 (G4 > G5 > G2 > G3), whereas the rank order for decreasing binding affinity (increasing 𝐾_𝑑_) follows the trend of G2 > G3 > G5 > G4 (Table 1). In the context of the maltodextrin transport system, which includes two orthogonal substrate-binding proteins, Athe_0598 could be considered analogous to Athe_2310 as the substrate-binding protein with higher specificity towards shorter oligosaccharides. Yet, unlike the maltodextrin system where one SBP does not display a significant affinity towards the disaccharide maltose, both Athe_0597 and Athe_0598 bind cellobiose as a cognate ligand, causing large melting temperature shifts (and in the case of Athe_0598, the only melting temperature shift) (22). The Glucan-Degradation-Locus (GDL) in *A. bescii,* encoded by the Athe_1860– Athe_1867 locus, is known to produce the CAZymes that primarily drive cellulose deconstruction (14,15,30). *A. bescii* has evolved to produce a native secretome that maximizes glucan conversion into cello-oligosaccharides (15,16). Prior results have shown that purified GDL enzyme cocktails or even the concentrated native secretome themselves are not as effective as an active *A. bescii* culture in degrading cellulose, reifying the significance of mass action kinetics such as cell adhesion to cellulose and simultaneous uptake of cello-oligosaccharides to remove end-product inhibition of GDL CAZymes and ensure continuous availability of cellobiose (15,16).

Multiple studies have suggested that the bioenergetic benefit of transporting higher-order cello-oligosaccharide substrates is vital for offsetting the significant metabolic costs of expressing a large array of extracellular CAZymes for cellulose deconstruction (35,36). The cost of translocating one sugar molecule across the membrane remains fixed at two ATP molecules regardless of sugar size; an affinity for tansporting larger sugars implies a more efficient use of resources. As cellobiose is the primary product of the synergistic activity of GDL CAZymes, the presence of a second highly transcribed ABC cellobiose-binding protein enables *A. bescii* to increase the rate of removing product inhibition of cellulases – and ensure that its transporter is always saturated with either short or long cello-oligosaccharides regardless of what is extracellularly available in their carbohydrate-limiting native environments.

Figure 1B shows that Athe_0597 is much more highly conserved across all *Caldicellulosiruptor* species compared to Athe_0598. Athe_0597 is also multi-functional, binding an array of cello-oligosaccharides while also binding cellobiose at association constants similar to Athe_0598. This further suggests that Athe_0597 plays the more important role in cellulose assimilation. And while all *Anaerocellum* and *Caldicellulosiruptor* species are hemicellulolytic, several *Caldicellulosiruptor* species including *C. kristjanssonii* and *C. hydrothermalis* are not cellulolytic (37). Yet even without the ability to hydrolyze cellulose, possessing this cello-oligosaccharide transporter locus likely continues to confer metabolic benefit by allowing scavenging of cello-oligosaccharides from the cellulolytic activity of other cellulolytic species in their native hot springs (37).

The inability of both strains HTAB187 (Δ*pyrE*Δ*athe_0595-0598*) and MACB1080 (Δ*pyrE*Δ*msmK*) to grow on cellulose strongly suggests that the promiscuous *msmK* (Athe_1803) is the ATPase associated with the Athe_0595–0598 cello-oligosaccharide transporter locus. This result is also consistent with prior MACB1080 growth performed on Avicel (13). Interestingly, while a previous study showed that MACB1080 can experience modest growth on cellobiose, we were unable to reproduce this result (13).

We suspect that possible variance in inoculation method between studies could account for this difference, or fructose carryover from the inoculum in this previous study facilitated the observed growth of MACB1080 in cellobiose-containing media (13,32).

HTAB187 demonstrated modest growth on cellobiose compared to the MACB1018 parent strain. Because HTAB187 on cellobiose attains final OD680 levels similar to growth on glucose, we speculate two possible causes for this phenomenon. First, it is possible that extracellular β-glucosidase activity, such as that of CelB, may be responsible for hydrolyzing cellobiose into glucose, the latter of which strain HTAB187 can still uptake. If this were true, however, we should also see partial MACB1080 growth on glucose enzymatic products in the media, which was not the case. Alternatively, *A. bescii* possesses other oligosaccharide transporters with some affinity for cellobiose uptake that still enable growth on the disaccharide. And because these redundant transporters may not possess as high a binding affinity for cellobiose, or because these transporters are not as highly expressed, the growth rate of HTAB187 on cellobiose is still severely impaired.

Cello-oligosaccharide uptake in a lignocellulolytic thermophile has also been studied in *Clostridium thermocellum* (32,38). Four transporters – transporter A, transporter B, transporter C, transporter D – were suggested to uptake cello-oligosaccharides based on ITC study of their respective substrate-binding proteins (33). While the substrate-binding protein (SBP) transporter A appears to only bind cellotriose, the respective SBPs of transporter B and transporter C bind cellobiose to cellopentaose (33). The SBP of transporter C also binds glucose, while the SBP of transporter D binds cellotriose to cellopentaose. At first glance, it would appear that Athe_0597 is analogous to transporter A in that it binds only a specific cello-oligosaccharide length (cellobiose in the case of Athe_0597 and cellotriose in the case of the SBP to transporter A), while the SBPs of transporters B, C, and D are analogous to Athe_0598 in their ability to bind both short and long cello-oligosaccharides. However, a more recent study incorporating both biophysics and genetics in *C. thermocellum* suggested that only transporter B mediates cello-oligosaccharide uptake (32). ITC characterization of the SBP to transporter A found that it only binds glucose; it is not apparent that the SBPs to transporters C and D bind cello-oligosaccharides of any length (32). This finding was unexpected at the time given that cellulose is a key growth substrate for *C. thermocellum* and that sugar transporters for primary carbon sources tend to be redundant in many microbes (32,39,40). The thermophilic bacterium *Thermotoga maritima* possesses three redundant maltodextrin transporters, while *A. bescii* is also known to possess two orthogonal maltodextrin transporters (41,42). Yet observing that *A. bescii* also possesses a single transporter for cellulose as a primary carbon source suggests that this physiological trait may not be as uncommon as originally thought.

It is most likely that HTAB187 fails to grow on cellulose because it cannot uptake the cello-oligosaccharides released. Transcriptomic and proteomic evidence suggests that expression of the GDL is not tied to uptake of a unique substrate, unlike cellobiose-dependent induction of the cellulosome in *C. thermocellum* (13,16,32). Decoupling the root cause of growth disruption on cellulose by observing simultaneous cellulose deconstruction alongside growth on non-cellulosic sugars is the most important next step.

In sum, this study identifies Athe_0595-0598 as the primary ABC transporter responsible for cello-oligosaccharide uptake in cellulolytic A. bescii. Our results demonstrate that Athe_0597 and Athe_0598 exhibit distinct binding affinities, with Athe_0597 binding a range of cello-oligosaccharides and Athe_0598 specializing in cellobiose uptake. Knockout of this transporter locus abolished cellulose utilization and severely impaired growth on cellobiose, underscoring its essential role in cello-oligosaccharide assimilation, a main carbon source for cellulolytic *A. bescii*. These findings not only advance our understanding of thermophilic sugar transport in the *Anaerocellum* and *Caldicellulosiruptor* genera, but also provide a foundation for engineering strategies aimed at optimizing *A. bescii* for lignocellulosic bioprocessing applications.

## Materials and Methods

### Bacteria & Growth Conditions

*E. coli* strains including NEB 5-α (New England Biolabs) and BL21 (DE3) pRosetta2 (EMD Millipore), were plated routinely on Luria-Bertani medium (10 g/L tryptone, 5 g/L yeast extract, 10 g/L NaCl) with 1.5% agar. All plates were supplemented with 50 μg/mL kanamycin for selection. All *E. coli* strains were cultured routinely in Luria-Bertani broth containing an additional 24 g/L yeast extract. All plates and cultures for growing *E. coli* BL21 (DE3) pRosetta2 strains were supplemented with 33 μg/mL chloramphenicol antibiotic. *A. bescii* strains DSMZ 6725, MACB1018, and MACB1080 were provided by the labs of Dr. Robert Kelly (North Carolina State University) and Dr. Michael W.W. Adams (University of Georgia) and cultured on complex media (C516) containing 5 g/L maltose, 0.5 g/L yeast extract, and 40 μM uracil as described previously (22). Late exponential phase *A. bescii* cells were harvested and pelleted by centrifugation at 5,000 x g for 20 min. Genomic DNA was extracted from pelleted cells using a Quick-DNA Miniprep kit (Zymo Research) and quantified using a UV-Vis Spectrophotometer Nanodrop (Thermo Scientific).

### Chemicals

The following monosaccharides and oligosaccharides were used in this study: D-Xylose (> 99%, Thermo Scientific), D-Glucose (Thermo Scientific), D-Cellobiose (> 98%, Acros Organics), D-Cellotriose (> 95%, Neogen Corporation), D-Cellotetraose (> 90%, Neogen Corporation), D-Cellopentaose (> 95%, Biosynth), Avicel® PH-101 Cellulose Resin (Sigma Aldrich).

### Substrate Binding Protein Cloning

Methods are performed as previously described in Tjo et al. 2024 (22). To recapitulate, genes encoding for the cello-oligosaccharide-binding proteins Athe_0597 and Athe_0598 absent their respective signal peptides (as predicted by SignalP 5.0) were amplified via PCR from *A. bescii* DSMZ 6725 genomic DNA. PCR amplification using Phusion polymerase (New England Biolabs) made use of primers described in Table S2 according to manufacturer instructions. Athe_0597 was inserted into a pRSF1-b backbone (gift from the Kelly Lab, North Carolina State University) whereas Athe_0598 was inserted into a pCri8a (Addgene Plasmid #61317) backbone. Gene insertion was performed via Gibson Assembly using the NEB HiFi Assembly Mastermix as detailed in the manufacturer instructions. N-terminal hexa-histidine-tags were added to each substrate-binding protein gene to enable protein purification via immobilized metal affinity chromatography (IMAC). The plasmid containing Athe_0597 is hereafter dubbed pHT007, while the plasmid containing Athe_0598 is dubbed pHT017. Each plasmid was transformed to *E. coli* NEB 5-α, with 50 μg/mL kanamycin selection pressure for plasmid maintenance. Plasmids were prepared and purified using the Zymo Research Plasmid Miniprep - Classic kit (Zymo Research) and sequenced via Plasmidsaurus’ whole plasmid sequencing service. Sequence confirmed pHT007 and pHT017 plasmids were each transformed into *E. coli* BL21 (DE3) pRosetta2 strains (EMD Millipore), with double selection markers of 50 μg/mL kanamycin and 33 μg/mL chloramphenicol for retention of both protein-containing and pRosetta2 vectors.

### Protein Expression & Purification

Methods are performed as previously described in Tjo et al. 2024 (22). *E. coli* BL21 (DE3) pRosetta2 strains transformed with respective pHT007 and pHT017 vectors were grown in 1 L ZYM-5052 auto-induction media supplemented with 50 μg/mL kanamycin and 33 μg/mL chloramphenicol in 2.8 L shake flasks at a temperature of 37 °C and shake speed of 250 rpm. Cell cultures were harvested after 20 hours of overnight growth at 4,500 x g for 20 min. Pellets were stored in a -20 °C freezer. IMAC Buffer A (20 mM sodium phosphate monobasic, 500 mM sodium chloride, pH 7.4) was used to resuspend cell pellets at a ratio of 10 mL buffer per 1 g of wet cell pellet. An Emulsiflex-C5 High-Pressure Homogenizer (Avestin) was used to lyse resuspended pellets via house air flow maintained at 30 psi. Each cell lysate was subjected to a pressure of approximately 100,000 kPa per pass and a minimum of two full lysis cycles. Homogenized cell lysates were then heat treated at a temperature of 68°C for 30 min using a water bath. Heat treated cell lysate was centrifuged using Nalgene® Oak Ridge High-Speed PPCO Centrifuge Tubes at 36,000 x g for 30 min (Thermo Scientific). The pooled supernatant was filtered through a 0.2 μm PES filter. Using a BioRad NGC 10 FPLC (Bio-Rad), filtered cell extract was fed into a 5 mL HisTrap HP Nickel-Sepharose (Cytiva) column in line with manufacturer specifications. Ni-NTA sample load, column wash, elution samples were selected via inspection of the absorbance curve at 280 nM in the chromatogram, and subsequently run using a precast 4-20% Mini-PROTEAN**®** TGX Stain-Free Protein Gel (Bio-Rad), using the Precision Plus Protein Standard (Bio-Rad) as the protein molecular weight ladder. Select purified elution fractions with high protein quantity and purity were pooled, concentrated, and buffer exchanged using a 10 kDa MWCO PES filter 20 mL Spin-X Concentrator (Corning). The final protein storage and characterization buffer is comprised of 50 mM HEPES, 300 mM NaCl, pH 7.0. A bicinchoninic acid (BCA) assay (Thermo Fisher Scientific) was used to quantity pHT007 concentrated to 50 mg/mL and pHT017 concentrated to 10 mg/mL. Both proteins were run on SDS-PAGE to verify protein identity by weight and sample purity (Figure S1).

### Isothermal Titration Calorimetry (ITC)

Methods are performed as previously described in Tjo et al. 2024 (22). Briefly, measurements were performed at a temperature of 25 °C using a MicroCal PEAQ Isothermal Titration Calorimeter (Malvern Panalytical) hosted by the Princeton University Biophysics Core Facility. Standard titration experiments were performed at a protein sample volume of 280 μL concentrated to 50 μM, as well as sugar ligand volume of 40 μL concentrated to 500 μM. A cell stir speed of 750 rpm was used throughout the entire titration experiment. Protein in the cell was initially injected with a priming aliquot of 0.2 μL sugar, succeeded by 19 injections at a volume of 2.0 μL each, with 2 min intervals between injections. A single site binding model was used to determine integrated heat effects via non-linear regression (Microcal PEAQ-ITC Analysis). Binding dissociation constant and key thermodynamic parameters were determined from the fitted isotherms via the Gibbs Free Energy form 𝛥𝐺 = −*RTln*(*K_d_*). The dissociation constant 𝐾_𝑑_ was calculated based on the binding isotherm’s slope at the equivalence point. The inverse of the 𝐾_𝑑_ is defined as the association rate constant 𝐾_𝑎_. 280 μL Milli-Q water was used as the reference.

### Differential Scanning Calorimetry (DSC)

Methods are performed as previously described in Tjo et al. 2024 (22). Briefly, melting curve profiles of hexa-histidine-tagged Athe_0597 and Athe_0598 were determined via DSC using the MicroCal PEAQ-DSC (Malvern Panalytical) hosted at the Princeton University Biophysics Core Facility. Athe_0597 and Athe_0598 were screened against a palette of β-glucans: cellobiose (G2), cellotriose (G3), cellotetraose (G4), and cellopentaose (G5). Glucose (G1) was also tested as a negative control. Each protein-sugar mixture contained final concentrations of 2 mg/mL protein and 5 mM sugar, with the same protein storage buffer of 50 mM HEPES, 300 mM NaCl, pH 7.0 used as the reference. All scan rates were set at 240 °C / min, reaching a maximum temperature of 130 °C. Raw data consisted of the heat capacity *C_p_* plotted as a function of temperature, followed by normalization by each respective run’s maximum heat capacity value, *C*_*p,max*_. *C_p,max_* is defined as *C*_*p,max*_ = *C_p_*(*T* = *T*_*m*_), where *T*_*m*_ is the melting temperature of the protein mixture. Each protein-sugar mixture was ascribed a 𝛥*T*_*m*_, defined as 𝛥*T*_*m*_ = *T*_*m,holo*_ − *T*_*mapo*_, where *T*_*m,holo*_ is the melting temperature of a given protein-sugar mixture and *T*_*m,apo*_ is the melting temperature of the protein in the absence of sugar.

### Construction of *A. bescii Δathe_0595-0598* Knockout Strain

Starting with the MACB1018 background parent strain (Δ*pyrE*), the *athe_0595-0598* locus was chromosomally deleted via homologous recombination using previously described methods (14,19) (Figure S2). The plasmid pKHT017 contained a ∼1 kb flanking region upstream of the Athe_0595 gene and a ∼1 kb flanking region downstream of the Athe_0598 gene; both flanking regions are adjacent one another in the vector (Figure S2). Flanking regions were PCR amplified from *A. bescii* DSMZ 6725 genomic DNA using Phusion polymerase (New England Biolabs), and inserted into a pGL103 backbone, as previously described (19). The knockout vector was transformed into *E. coli* NEB 5-α via heat shock at a temperature of 42 °C, subsequent recovery in SOC outgrowth medium (New England Biolabs), and grown on plates and cultures supplemented with 50 μg/mL kanamycin and 50 μg/mL apramycin selection pressures for plasmid maintenance. pKHT017 was prepared and purified using the Zymo Research Plasmid Miniprep - Classic kit (Zymo Research) and sequenced via Plasmidsaurus’ whole plasmid sequencing service as described above. Sequence confirmed pKHT017 transformants were grown in 250 mL Luria-Bertani medium supplemented with additional 24 g/L yeast extract, as well as 50 μg/mL kanamycin and 33 μg/mL apramycin antibiotics, in 1 L shake flasks at a temperature of 37 °C. Incubator shake speed was set at 250 rpm. Cells were pelleted and processed via a ZymoPURE™ II Plasmid Purification Maxiprep kit. pKHT017 plasmid was then methylated *in vitro* using M.CbeI methyltransferase as previously described (19,43). Successful methylation was verified by protection against HaeIII restriction endonuclease activity.

Competent MACB1018 cells were prepared in Low-Osmolarity-Defined (LOD) media containing 5 g/L cellobiose and 40 μM uracil, as described previously (13,19). LOD media and all media hereafter described were degassed and made anaerobic for 15 minutes using a vacuum pump, with a gaseous headspace comprising 80% (v/v) nitrogen and 20% (v/v) carbon dioxide gas. Competent cells were electroporated using a Bio-Rad Gene Puler with 2 μg of methylated pKHT017 plasmid at 2.2 kV, resistance of 400 𝛺, and capacitance of 25 μF. 1 mL of recovery media, containing modified DSM 516 media supplemented with 5 g/L of yeast extract, and pre-heated to a temperature of 70 °C, was immediately used to resuspend electroshocked cells (13). After both 60 min and 120 min post-electroporation, 5 mL of electroporated cells in recovery medium were transferred to 50 mL (selective) DSM 516 medium supplemented with 50 μg/mL kanamycin. Cloudy media after 48 - 72 hours of growth at 70 °C suggested successful transformations; 1 mL samples of such cultures were passaged into 10 mL fresh selective DSM 516 media containing 50 μg/mL kanamycin. Passaged transformants were then plated in solid, selective modified DSM 516 medium with 1.5% agar, grown for 48 - 72 hours at 70 °C under 95% (v/v) nitrogen and 5% (v/v) hydrogen gas, and single colonies selected and screened via PCR to confirm knockout vector chromosomal integration adjacent to the Athe_0595 - 0598 locus (13). A PCR-verified first crossover colony is then plated on solid, modified DSM 516 medium containing 8 mM 5-fluoroorotic acid (5-FOA) and 40 μM uracil. The 5-FOA counter-selection step also marks a permanent change in growth substrate from 5 g/L cellobiose to 5 g/L maltose. Single colonies were picked and screened for complete deletion of the Athe_0595 - 0598 locus following counter-selection via colony PCR using primers directly outside the 1 kb upstream and downstream flanking regions used in the pKHT017 vector.

### *A. bescii* Growth Curves on Select Carbohydrate Substrates

Freezer stocks of MACB1018 (Δ*pyrE*), MACB1080 (Δ*pyrE*Δ*msmK*), and HTAB187 (Δ*pyrE*Δ*athe_0595- 0598*) were each thawed and used to inoculate 50 mL of anaerobic, non-selective modified DSM 516 medium containing 40 μM uracil and 5 g/L of fructose in 100 mL serum bottles for overnight growth. Strains were passaged once more into anaerobic, non- selective modified DSM 516 medium containing 40 μM uracil and 5 g/L of fructose. Upon reaching late exponential phase at approximately 16 hours of growth at 70 °C, cultures were cooled to room temperature. 10 mL of cells from each strain were extracted and centrifuged at 14,000 rpm for 2 min. The supernatant was primarily decanted via pouring out the liquid, with residual supernatant removed via pipetting. The pellet was then washed and resuspended in 1 mL of non-selective modified DSM 516 sugar-free medium, and then injected into 10 mL of the same sugar-free medium contained within a 18 x 125 mm anaerobic tube sealed by 20 mm butyl rubber stoppers (Duran Wheaton Kimble), resulting in a sugar-free bacterial culture with final 𝑂𝐷_680_ ∼ 0.10.

1 mL of the sugar-free bacterial culture was used to inoculate 50 mL of non- selective modified DSM 516 medium containing 40 μM uracil and a single carbohydrate source at a density of 5 g/L sealed by 20 mm butyl rubber stoppers (Duran Wheaton Kimble). All inoculated cultures reported an initial 𝑂𝐷_680_ ∼ 0.01, and were grown at a temperature of 70 °C, without shaking, for 40 hours in biological triplicate, as previously described (13). The 𝑂𝐷_680_ of growth cultures were measured at intervals of roughly 2.5 - 6.0 hours using the cuvette setting of a UV-Vis Nanodrop Spectrophotometer, with 1X DSM 516 salt solution as the blank. Cell protein content as a proxy for cell density under Avicel growth conditions was quantified using the Bradford method with bovine serum albumin (BSA) protein standards (Pierce Thermo Scientific) in a 96-well plate format, as previously described (13,32,44).

### Bioinformatics

Through PDB sequence similarity search, glucan-binding protein orthologs with solved crystal structures were identified using BLAST against 0597 and 0598 as query protein sequences (45). For Athe_0597, structures were from *Streptococcus pneumoniae* (PDB: 5SUO, 5SWA, 5SWB). For Athe_0598, structures included *Streptococcus pneumoniae* (PDB: 5SWA) as well as *Caldanaerobius polysaccharolyticus* (PDB: 4G68)*, Paenibacillus sp. str. FPU-7* (PDB: 7EHP), and *Thermotoga maritima* (PDB: 2O7I) (34,46–48). Amino acid sequences of all homologs used in this study are summarized in Table S3. PROMALS3D and ESPript 3.0 visualization software was used to identify similarities in secondary and tertiary protein structures from structure-based sequence alignment of each CBP and their orthologs (49,50).

### Ligand Docking Models

ColabFold v1.5.5 supplied with AlphaFold 2.0 parameters were used to generate the initial models; the best-match homolog was used as the template, with other homologs as the MSA (51,52). The “ref2015” score function under the Rosetta *relax* protocol was used to minimize AlphaFold-generated structures. Docking of carbohydrate ligands onto protein binding pockets was performed using the ROSIE server (53,54). Initial posing of the carbohydrate ligand within the protein binding pocket was based on homology to the corresponding crystal structure. 200 conformers of each carbohydrate substrate were generated using BCL (55). Each protein–substrate combination yielded 200 docked structures, and interface energies of binding were quantified from the ten lowest energy structures. KDEEP, a 3D convolutional neural network method for assessing interface energy, was used to further evaluate ligand-bound pose (56). All protein structures were visualized using PyMOL (57).

## Acknowledgements

This work was supported by a Roberto Rocca Graduate Fellowship from Techint Group to H.T.; by the High Meadows Environmental Institute at Princeton University through the generous support of the William Clay Ford, Jr ‘79 and Lisa Vanderzee Ford ‘82 Graduate Fellowship Fund to H.T.; by a National Science Foundation Graduate Research Fellowship DGE-2039656 to V.J.; by a Gordon Wu Fellowship to V.J.; by startup funds from the Department of Chemical and Biological Engineering and the Omenn-Darling Bioengineering Institute at Princeton University to J.A.J.; by seed funds from the School of Engineering and Applied Science and startup funds from the Department of Chemical and Biological Engineering at Princeton University to J.M.C.; all authors acknowledge financial support from Princeton University’s Omenn-Darling Bioengineering Institute. We acknowledge Venu Gopal Vandavasi, Director of the Princeton Biophysics Core, for instrument training and advice on experimental design. We also acknowledge Tom Silhavy, Ned Wingreen, and A. James Link for helpful discussions.

